# Base exchange inhibitors of SARM1 form mononucleotide adducts and activate SARM1 in vivo

**DOI:** 10.1101/2025.04.22.649875

**Authors:** Holly T. Reardon, Carlos Guijas, Jason R. Clapper, Julia A. Callender, Ellen M. Kozina, Francisco J. Sarabia, Clayton Hutton, Dylan M. Herbst, Taylor Andalis, Luka Perehinets, Cassandra L. Henry, Terry P. Lebold, David A. Kummer, Jacqueline L. Blankman, John J.M. Wiener, Klaus B. Simonsen, Tarek A. Samad

## Abstract

Activation of the NAD hydrolase SARM1 causes neuronal pathology. NAD binding to an allosteric site maintains SARM1 autoinhibition while NMN binding enables NADase activity. We evaluated SARM1 base exchange inhibitors (BEI) that exchange with nicotinamide in the SARM1 catalytic site to form inhibitor adducts. We found that BEI paradoxically activated SARM1 in some contexts, elevating both the SARM1 product cADPR and ratio to substrate, cADPR/NAD, in uninjured sciatic nerve and in healthy animals. Catalytically inactive E642A SARM1 knock-in mice were protected from elevated cADPR and cADPR/NAD in sciatic nerve after BEI dosing. Like the SARM1 activating forms of the neurotoxins Vacor and 3-acetylpyridine, BEI formed mononucleotide (MN) adducts in vivo. BEI also formed MN adducts in human THP-1 cells, as did the pyridine-containing drugs indinavir and chloroquine. These findings underscore the promiscuity of nicotinamide-related metabolism and potential for neurotoxicity arising from SARM1’s susceptibility to activation by exogenous agents.

## Introduction

The enzyme Sterile alpha and TIR motif containing 1 (SARM1) plays a key role in axon degeneration, and mice lacking SARM1 are protected in a wide variety of peripheral neuropathy, neurodegeneration, and nerve injury models (1, 2). SARM1 is normally autoinhibited in healthy neurons and becomes activated in disease and nerve injury contexts (3, 4). When active, SARM1 hydrolyzes nicotinamide adenine dinucleotide (NAD^+^, referred to here as NAD) to generate products such as cyclic adenosine 5’-diphosphate ribose (cADPR), adenosine diphosphate ribose (ADPR), and nicotinamide (5). Activation of SARM1’s NADase activity results in rapid loss of NAD combined with calcium mobilization through cADPR signaling, initiating a cascade of events culminating in physical destruction of axons and/or neurons (6). SARM1 is well established as the final executioner of the distinctive “dying back” distal axonal pathology known as Wallerian degeneration (7), but neuronal cell death can also occur when SARM1 becomes activated throughout the cell (8, 9).

Structural studies have led to greater understanding of mechanisms regulating SARM1 activation. SARM1 protein contains an N-terminal autoinhibitory armadillo repeat (ARM) domain, tandem sterile-alpha motif (SAM) domains that enable protein monomers to oligomerize into an octameric complex, and a C-terminal catalytic Toll/interleukin-1 receptor (TIR) domain (10). SARM1’s NADase activity is regulated by either NAD or nicotinamide mononucleotide (NMN) binding to an allosteric site within the ARM domain (4). NAD binding maintains the inactive conformation while NMN binding triggers a conformational change that dimerizes TIR domains to form the functional catalytic site (3, 4, 11).

Due to the opposing effects of NAD and NMN competing for the same allosteric regulatory site, the activation state of SARM1 is ultimately determined by the ratio of NMN/NAD in the surrounding microenvironment, with SARM1 normally held in the autoinhibited state in healthy neurons in which the concentration of NAD far exceeds NMN (12). Various disease states, nerve injuries, and toxic insults compromise NAD synthesis from the precursor NMN, resulting in excess NMN that displaces NAD from the allosteric regulatory site and relieves autoinhibition. The promiscuity of this allosteric mechanism makes SARM1 susceptible to activation by exogenous compounds; for example, both the former rodenticide Vacor and the neurotoxin 3-acetylpyridine (3-AP) are metabolized to form mononucleotide (MN) adducts that mimic the effect of NMN and activate SARM1, accounting for the neurotoxic effects of these agents (Figure 1A) (8, 9).

**Figure 1.**
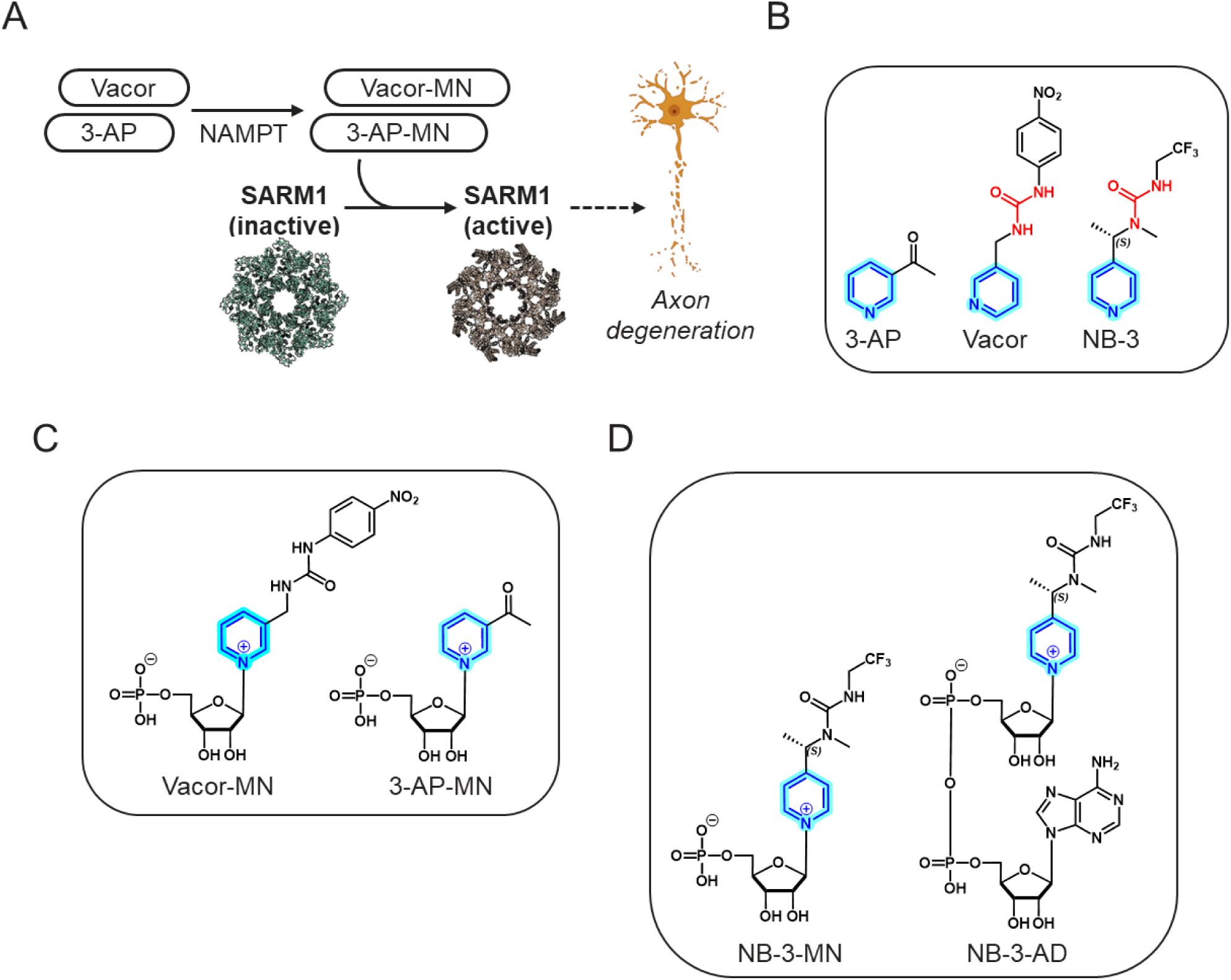
Mechanism for SARM1 activation by exogenous neurotoxins. **(A)** Vacor and 3-acetylpyridine (3-AP) are metabolized to form mononucleotide (MN) adducts that mimic the effect of NMN and activate SARM1, accounting for the neurotoxic effects of these agents. Structures for inactive SARM1 (green; PDB 7CM2) and active SARM1 (orange; PDB 7NAL1) are shown. Created with Biorender.com. **(B)** Structures of 3-acetylpyridine (3-AP), Vacor, and the base exchange inhibitor NB-3. **(C)** Known mononucleotide adducts associated with neurotoxicity. **(D)** Mononucleotide and dinucleotide adducts formed by NB-3.

Human genetic evidence underscores the deleterious effects of excessive SARM1 activation. For example, rare SARM1 genetic variants with loss of autoinhibition and increased constitutive NADase activity are enriched in sporadic amyotrophic lateral sclerosis (ALS) patients; the most common variant, V184G, causes motor neuron dysfunction when introduced into mice (13, 14). Loss-of-function mutations in the NMNAT (nicotinamide mononucleotide adenylyltransferase) enzymes, which catalyze the conversion of NMN to NAD, lead to an increase in the NMN/NAD ratio and concomitant SARM1 activation; loss of NMNAT2 leads to childhood onset progressive polyneuropathies (6, 15–18), while mutations in NMNAT1 (cytoplasmic only in retina) lead to congenital loss of photoreceptors in affected patients (19–23).

Thus, preventing SARM1 activation or inhibiting the activated NADase may be promising therapeutic strategies for neurological disorders. One approach to inhibition hijacks a natural base exchange reaction mechanism that can occur with free pyridines when NAD is bound in SARM1’s catalytic site; for example, activated SARM1 can exchange nicotinic acid for nicotinamide in NAD to generate nicotinic acid adenine dinucleotide (NAAD) as a side product (24). Several groups have designed compounds that readily exchange for nicotinamide in NAD, generating slow off-rate adenine dinucleotide (AD) adducts that act as orthosteric inhibitors of SARM1’s NADase activity (25–27). When we tested base exchange inhibitors (BEI) in vivo, we were surprised to observe paradoxical SARM1 activation in uninjured nerves. Intrigued by the structural similarity of BEI with the SARM1 activators Vacor and 3-AP (Figure 1B), we investigated whether BEI may activate SARM1 through a mechanism analogous to the MN adducts of Vacor and 3-AP (Figure 1C). As reported here, in addition to the expected BEI-AD adducts we also confirmed the presence of BEI-MN adducts (Figure 1D) correlated with SARM1 activation and characterized mechanisms of adduct formation in different tissue compartments, as well as broader potential for MN adduct formation with pyridine-containing approved drugs.

## Results

### BEIs elevate SARM1 activity biomarkers in uninjured nerve

Aiming to replicate reported neuroprotective effects of BEI in vivo (27), we tested compound NB-2 in a unilateral sciatic nerve axotomy (SNA) model (Figure 2A). In this model, axotomy triggers Wallerian degeneration of distal sciatic nerve, which can be assessed with the neurodegeneration biomarker neurofilament light chain (Nf-L). Oral administration of 30 and 100 mg/kg NB-2 at the time of axotomy strongly suppressed elevation of plasma Nf-L at 20 h post-axotomy (Figure 2B), consistent with a neuroprotective effect in the axotomized nerve. To confirm that these effects were due to SARM1 inhibition, we also measured SARM1’s product, cADPR, which was reported to be primarily generated by SARM1 activity in neuronal tissues (5). In distal sciatic nerve ipsilateral to the axotomy, the axotomy-induced rise in cADPR was reduced in 100 mg/kg NB-2-dosed animals compared to vehicle controls (Figure 2C). To confirm these results, we normalized cADPR to NAD in sciatic nerve as a measure of SARM1’s product to substrate ratio and observed dose-dependent reductions in cADPR/NAD in the injured ipsilateral nerve (Figure 2D). As a negative control, we also examined cADPR and cADPR/NAD in uninjured contralateral nerves from axotomized animals. We were surprised to detect a small but consistent elevation of both cADPR and cADPR/NAD in uninjured contralateral nerves in NB-2-dosed animals relative to vehicle controls, both when NB-2 was administered at the time of axotomy (Figure 2E) or at 30 or 100 mg/kg 8 h post-axotomy (Figure 2F). This effect was not limited to NB-2; the related compound NB-3 (27) and a BEI from a different chemical series identified in internal screening, A1715CG, were dosed at 100 mg/kg in the same model 8 h post-axotomy and also resulted in elevated cADPR and cADPR/NAD in the uninjured side (Figures 2G, H).

**Figure 2.**
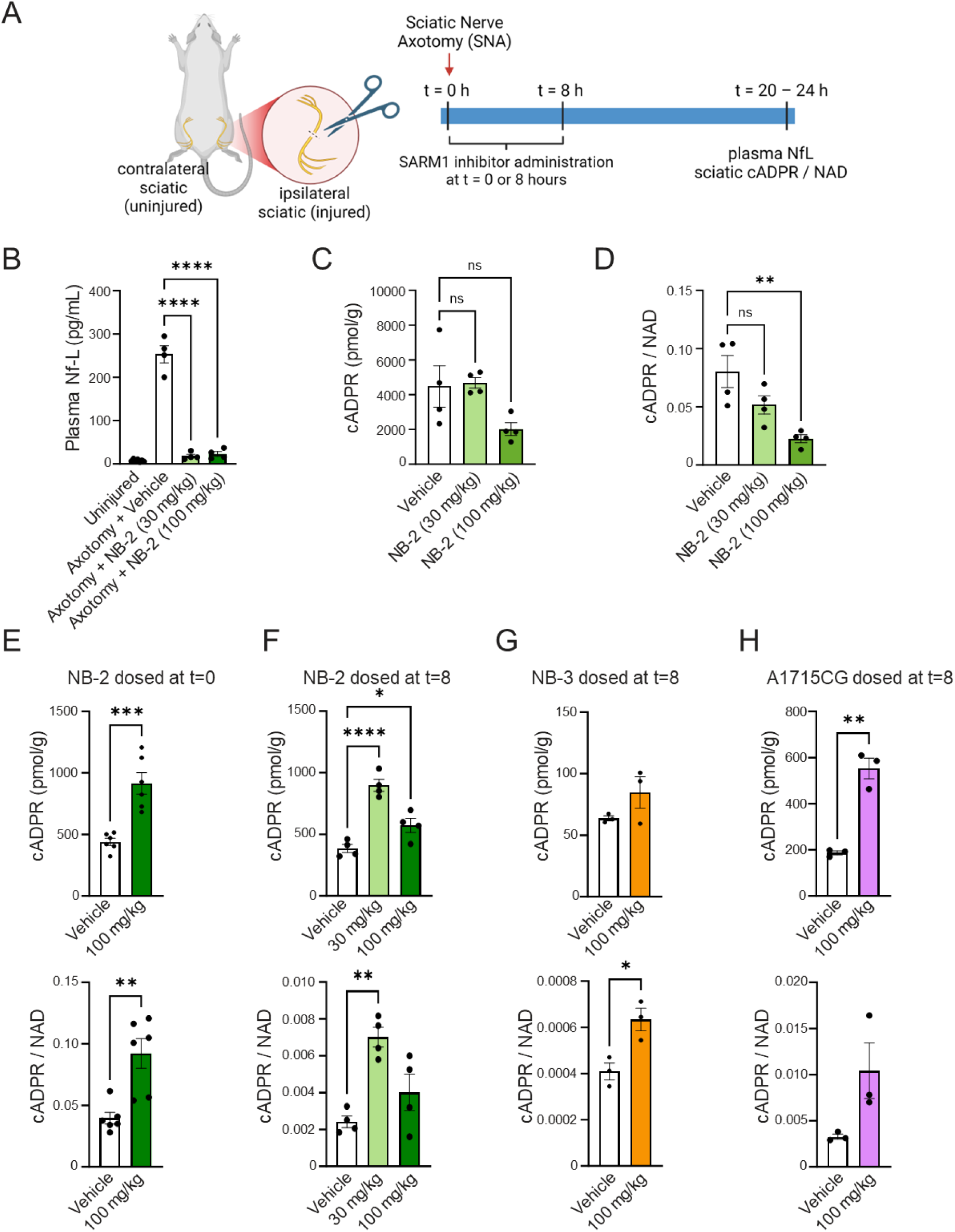
Base exchange inhibitors activate SARM1 in uninjured sciatic nerve. **(A)** Schematic depicting the study design to measure SARM1 activation during a unilateral sciatic nerve axotomy in rats. Created with Biorender.com. **(B)** Plasma Nf-L measurements 24 h after axotomy in animals dosed with NB-2 at the time of axotomy. **(C -D)** Ipsilaterial (injured) sciatic nerve cADPR alone **(C)** or normalized to NAD **(D)** 24 h after axotomy in animals dosed with NB-2 at the time of axotomy. **(E)** Contralateral (uninjured) sciatic nerve cADPR and cADPR/NAD ratio 24 h after axotomy, with 100 mg/kg NB-2 (green) dosed at the time of axotomy. **(F-H)** Contralateral (uninjured) sciatic nerve cADPR and cADPR/NAD ratio 20 h after axotomy, with 100 mg/kg NB-2 (green, **F**), 100 mg/kg NB-3 (orange, **G**) or 100 mg/kg A1715CG (purple, **H**) dosed 8 h after axotomy. Results are shown as mean ± SEM. N ≥ 3 animals per condition. *, p<0.05; **, p<0.01; ***, p<0.001, ****, p<0.0001 calculated using One-way ANOVA with Tukey’s multiple comparison test.

### BEIs form an MN adduct in vivo

Based on structural similarity to known SARM1 activators vacor and 3-AP, we hypothesized that BEI may also cause SARM1 activation through the formation of mononucleotide (MN) adducts. To test this hypothesis, we analyzed sciatic nerve samples from SNA studies where elevated cADPR/NAD had been observed. In addition to parent BEI compound and the previously described BEI-AD adducts known to be the active form of inhibitor (27), we also detected a BEI-mononucleotide adduct (MN) in both axotomized and uninjured sciatic nerves of animals dosed with BEI. Levels of NB-2-MN adducts increased along with dose of parent NB-2 compound (30 and 100 mg/kg, Figure 3A). All chemotypes of BEI that caused elevated cADPR/NAD also formed MN adducts in sciatic nerve, including 100 mg/kg NB-3 (Figure 3B) and 100 mg/kg A1715CG (Figure 3C). Interestingly, while for all BEI much higher levels of AD adduct were detected in injured ipsilateral nerves, MN adducts were higher in ipsilateral nerve for two compounds (NB-2 and A1715CG) while for NB-3 levels of MN adducts were not significantly different in ipsilateral compared to contralateral nerve. Neither MN nor AD adducts were detectable in plasma (data not shown).

**Figure 3.**
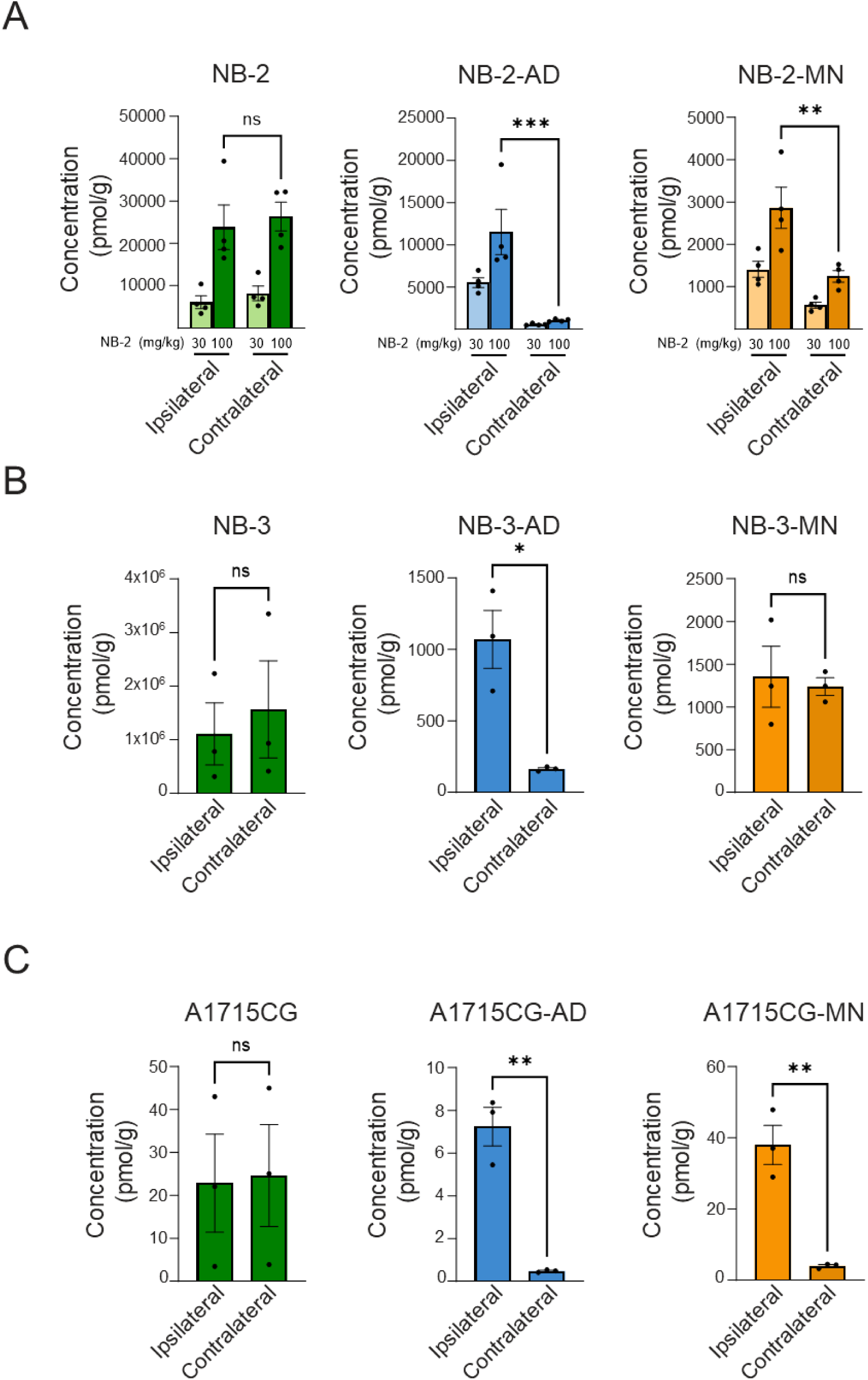
Base exchange inhibitors form a mononucleotide adduct in sciatic nerve. Depicted are the parent compounds (green), dinucleotide adducts (blue), and mononucleotide adducts (orange) detected in sciatic nerves from SNA studies with BEI. **(A)** NB-2 and its dinucleotide (NB-2-AD) and mononucleotide (NB-2-MN) adducts measured in sciatic nerve samples (ipsilateral and contralateral) from SNA studies 24 h after axotomy in rats dosed with 30 mg/kg or 100 mg/kg NB-2 at the time of axotomy. **(B-C)** Adducts measured in sciatic nerve 20 h after axotomy in rats dosed 8 h after axotomy with 100 mg/kg NB-3 **(B)**, or 100 mg/kg A1715CG **(C)**. Results are shown as mean ± SEM. N ≥ 3 animals per condition. *, p<0.05; **, p<0.01; ***, p<0.001, calculated using One-way ANOVA with Tukey’s multiple comparison test.

### MN adduct formation correlated with SARM1 activity biomarkers

To further investigate the correlation of SARM1 activation with MN adduct formation, we studied the kinetics of adduct formation and elevated cADPR/NAD using the BEI NB-2. Following a single oral dose of 30 mg/kg NB-2 in naïve rats, NB-2 adducts showed an accumulating trend in sciatic nerve from 4 to 24 h after dosing, while levels of parent compound dropped rapidly in both sciatic nerve and plasma over 24 h (Figure 4A). In the same study, levels of cADPR/NAD in sciatic nerve increased by 1.8-fold from 4 to 24 h in dosed animals (Figure 4B).

**Figure 4.**
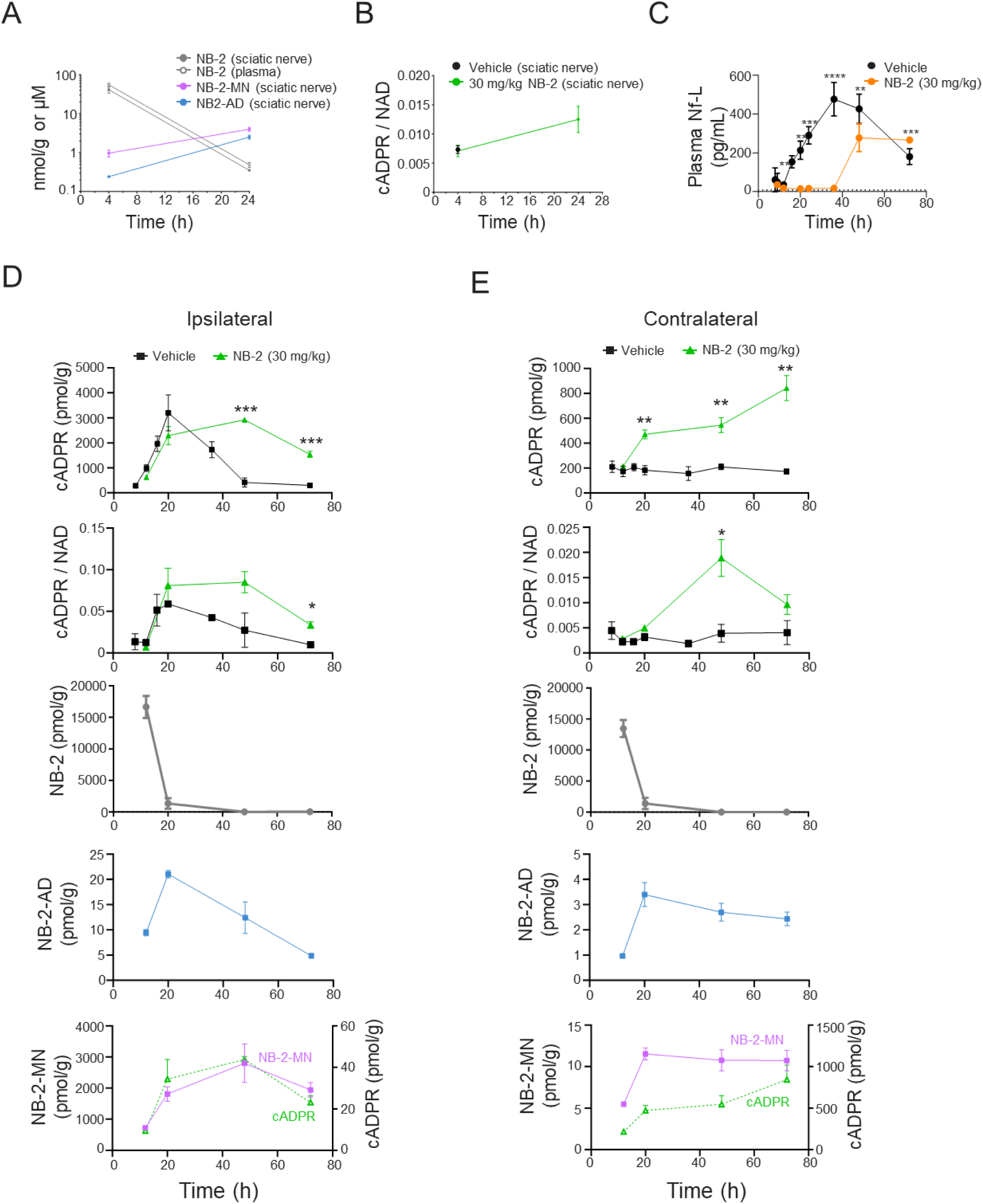
Mononucleotide adduct formation is correlated with SARM1 activation. **(A-B)** Nucleotide adducts **(A)** and cADPR/NAD **(B)** were measured in sciatic nerve from naïve animals dosed with NB-2 for 4 or 24 h. **(C)** Plasma NfL in rats treated with NB-2 (30 mg/kg) 8 h after sciatic nerve axotomy was measured at time points shown. **(D-E)** Time course of ipsilateral (injured, **D**) or contralateral **(E)** sciatic nerve cADPR, cADPR/NAD, NB-2, and NB-2 nucleotides in rats treated with NB-2 (30 mg/kg) 8 h after sciatic nerve axotomy (same study depicted in **C**). Results are shown as mean ± SEM. N ≥ 3 animals per condition. *, p<0.05; **, p<0.01; ***, p<0.001, calculated using an unpaired t-test.

A more detailed time course of NB-2 was evaluated in the SNA model. A single dose of NB-2 (30 mg/kg) was administered 8 h after axotomy, followed by measurement of plasma Nf-L and levels of cADPR, cADPR/NAD, NB-2, NB-2-AD, and NB-2-MN in both ipsilateral and contralateral sciatic nerves at time points up to 72 h post-axotomy. NB-2 dosing resulted in complete suppression of plasma Nf-L elevation for 36 h, followed by a period of increasing Nf-L ending with significantly elevated Nf-L relative to vehicle control at 72 h (Figure 4C). In contrast, vehicle treated animals exhibited peak Nf-L at 36 h followed by consistently declining Nf-L levels. In ipsilateral nerve (Figure 4D), maximal suppression of SARM1 activity biomarkers was observed at 20 h post-axotomy in NB-2 dosed animals, with a 28% reduction in sciatic nerve cADPR but no apparent change in cADPR/NAD. However, NB-2 dosed animals exhibited significantly elevated cADPR at 48 and 72 h relative to vehicle control animals, with a similar trend in cADPR/NAD. While NB-2 levels in ipsilateral nerve dropped rapidly over the first 20 h, the highest concentrations of AD adduct were at 20 h post-axotomy, corresponding with maximal reductions in cADPR and cADPR/NAD. However, as levels of NB-2-AD adduct were dropping at 48 h, levels of NB-2-MN continued to increase in ipsilateral sciatic nerve. Strikingly, the increase in ipsilateral NB-2-MN at 48 h corresponded with a peak in both ipsilateral cADPR and cADPR/NAD at the same time point. By 72 h, levels of both the MN adduct and cADPR/NAD had fallen, although cADPR/NAD was still highly elevated compared to vehicle control animals. In uninjured contralateral sciatic nerve (Figure 4E), elevated cADPR was observed in NB-2 dosed animals as early as 20 h post-axotomy and continued to be significantly elevated at 48 and 72 h. Peak levels of cADPR/NAD were observed at 48 h in contralateral nerve. Levels of both adducts in the uninjured nerve increased while NB-2 levels fell during the first 20 h. NB-2-AD levels peaked at 20 h but did not decrease as rapidly as in ipsilateral nerve. NB-2-MN levels in contralateral nerve remained two-fold higher at 48 and 72 h compared to 12 h, again mirroring the continued elevation in cADPR and cADPR/NAD at 48 and 72 h.

### BEI activate SARM1 in healthy mice

To understand these effects, it was important to verify that the elevation in cADPR/NAD we observed in both injured and healthy sciatic nerve after BEI dosing was due to SARM1 activation. To this end, we used catalytically-inactive SARM1 CRISPR knock-in (KI) mice with the catalytic glutamate of SARM1 mutated to alanine (E642A). We had previously demonstrated that these mice were completely protected from elevated Nf-L and cADPR/NAD in the SNA model, confirming that they lacked SARM1 activity and that the biomarkers measured in SNA were solely attributed to SARM1 (Supplemental Figure S1). Dosing the BEI NB-3 in naïve animals (no axotomy on either side) at 100 mg/kg resulted in elevated sciatic nerve cADPR/NAD 24 h after administration in wild type (wt, SARM1^+/+^) and heterozygous (SARM1^+/E642A^) animals relative to vehicle controls but had no effect on sciatic cADPR/NAD in homozygous SARM1 KI (SARM1^E642A/E642A^) mice (Figure 5A), confirming that the elevated cADPR/NAD was due to SARM1 activity.

**Figure 5.**
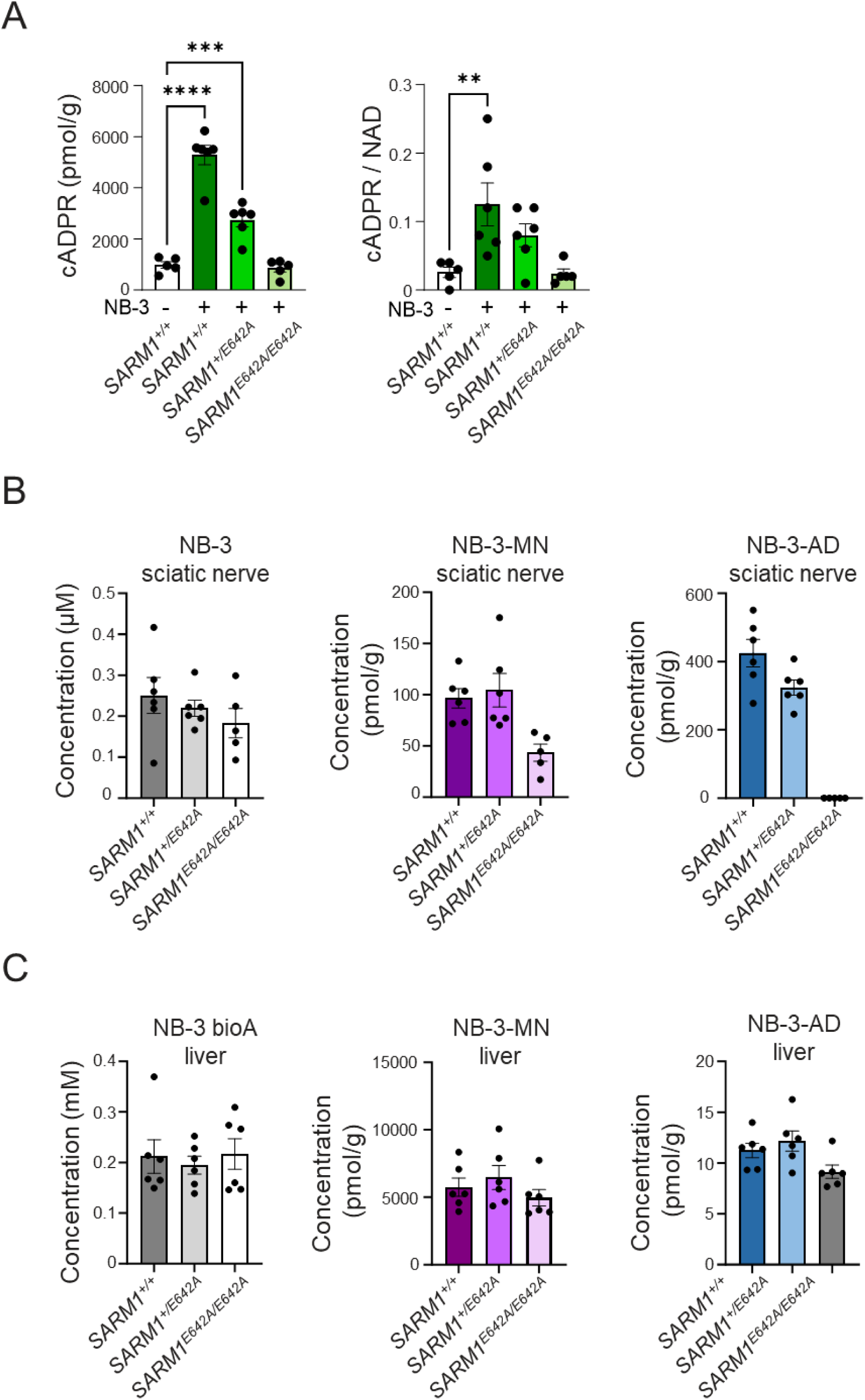
Base exchange inhibitors activate SARM1 in naïve animals. **(A)** cADPR and cADPR/NAD ratio after dosing 100 mg/kg NB-3 in naïve CRISPR knock-in mice carrying a catalytically-inactive SARM1 mutation (E642A). **(B)** NB-3 parent, -MN, or -AD adducts were measured in sciatic nerves of naïve *SARM1^E642A^* animals. **(C)** NB-3 parent, -MN, or -AD adducts were measured in livers of naïve *SARM1^E642A^* animals. Results are shown as mean ± SEM. N ≥ 5 animals per condition. **, p<0.01; ***, p<0.001, ****, p<0.0001 calculated using One-way ANOVA with Tukey’s multiple comparison test.

### Adducts form in multiple tissues through multiple enzymes

Since BEI-AD adducts have been proposed to form in the active site of SARM1 (25, 27), we wondered if SARM1activity was also contributing to formation of MN adducts. To understand the role of SARM1, we administered 100 mg/kg NB-3 to naïve SARM1 KI mice and wt controls, and measured levels of NB-3, NB-3-MN, and NB-3-AD in sciatic nerve and liver 24 h after dosing. In sciatic nerves (Figure 5B) we observed NB-3-MN adduct in wt and heterozygous SARM1 KI mice, but NB-3-MN levels in homozygous SARM KI sciatic nerves were approximately 50% lower than levels in wt, suggesting that MN adducts in sciatic nerve occurred partly through SARM1 activity and partly through some other mechanism. The NB-3-AD adduct was detected in wt and heterozygous SARM1 KI sciatic nerves and not in homozygous SARM1 KI, as expected from previous reports (25, 27). In liver (Figure 5C), NB-3-MN adducts were detected at similar levels in all genotypes, confirming that liver MN adduct formation was entirely independent of SARM1. Interestingly, levels of NB-3-MN adducts were much higher in liver than in sciatic nerve. We were also surprised to discover NB-3-AD was present in liver in all genotypes, suggesting that in liver a SARM1-independent mechanism exists to generate the inhibitor adduct.

### Human cells form BEI-MN adducts

Since all our results thus far were in rodents, we decided to test whether adduct formation could occur in a human cell line. It has been reported that vacor-MN is formed through the activity of the enzyme nicotinamide phosphoribosyltransferase (NAMPT) (9, 28), so we selected the human THP-1 cell line, which has been used as a model system for optimization of NAMPT inhibitors (29). Drawing from literature (27) and internal screening, we assembled a panel of BEI that exhibited elevated cADPR in contralateral sciatic nerve when dosed in the SNA model at 100 mg/kg (Table 1). We found that BEI incubated overnight in THP-1 at 100 µM (comparable to peak unbound plasma concentrations achieved in vivo with efficacious doses of BEI (27)) reprised the formation of MN adducts that we had observed in rodent sciatic nerve after in vivo dosing. Although sciatic nerve samples were not available to test for in vivo MN adducts with all dosed compounds, we confirmed that BEI-MN adducts formed in THP-1 cells for all BEI chemotypes associated with elevated cADPR in uninjured rodent sciatic nerve. In contrast, the BEI-AD adducts formed in vivo were not always detected in THP-1 cells; for example, no AD adduct was detected in THP-1 for the enantiomer of NB-3, A1769LH, despite detection in sciatic nerve from SNA experiments. Interestingly, A1769LH robustly formed MN adducts both in vitro and in vivo.

**Table 1:**
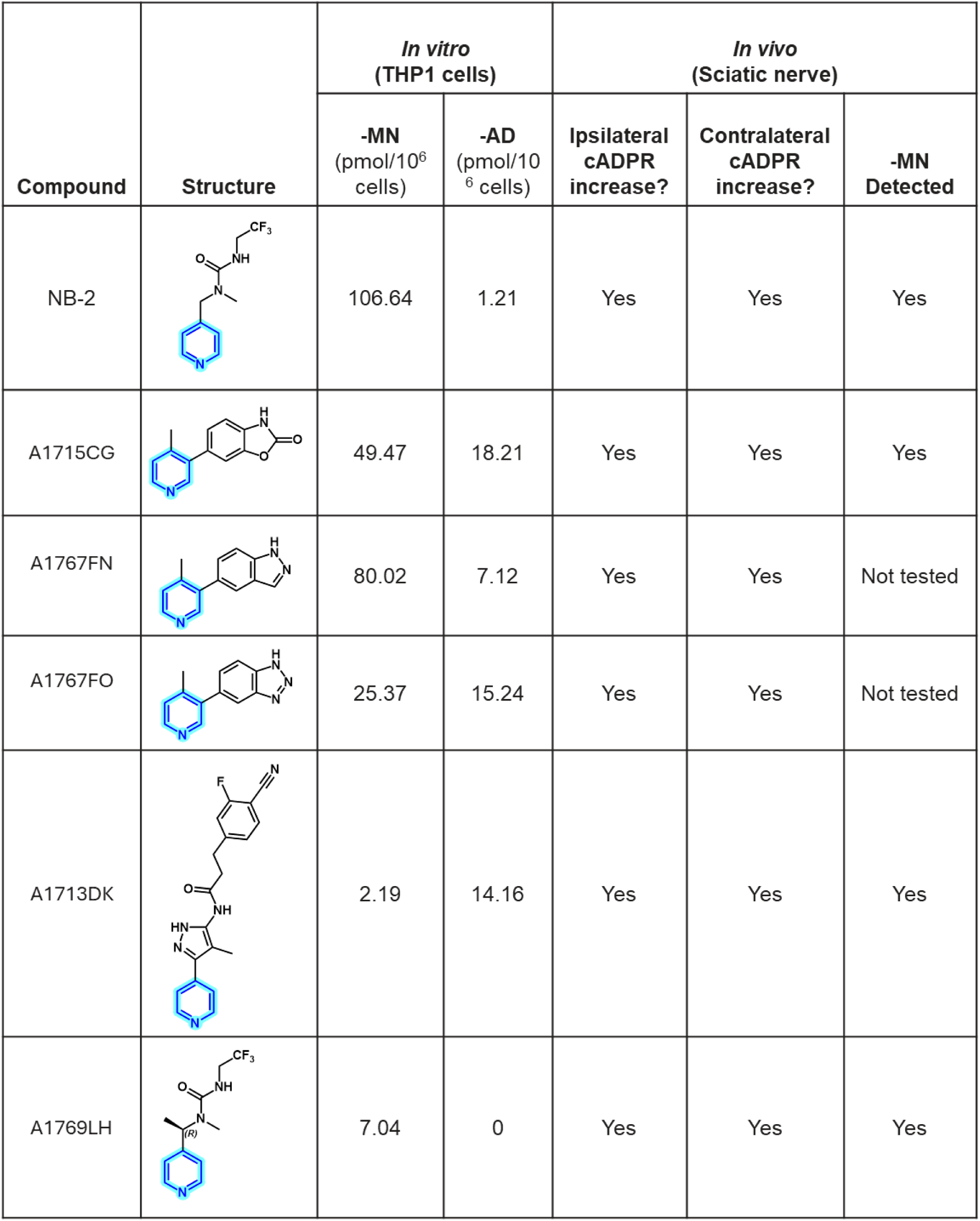
Mononucleotideadducts form in human TMP-1 cells *in vitro* and in sciatic nerve *in vivo*.

| Compound | Structure | *In vitro*<br>(THP1 cells) |  | *In vivo*<br>(Sciatic nerve) |  |  |
| --- | --- | --- | --- | --- | --- | --- |
|  |  | -MN<br>(pmol/10<sup>6</sup><br>cells) | -AD<br>(pmol/10<sup>6</sup><br>cells) | Ipsilateral<br>cADPR<br>increase? | Contralateral<br>cADPR<br>increase? | -MN<br>Detected |
| NB-2 |  | 106.64 | 1.21 | Yes | Yes | Yes |
| A1715CG |  | 49.47 | 18.21 | Yes | Yes | Yes |
| A1767FN |  | 80.02 | 7.12 | Yes | Yes | Not tested |
| A1767FO |  | 25.37 | 15.24 | Yes | Yes | Not tested |
| A1713DK |  | 2.19 | 14.16 | Yes | Yes | Yes |
| A1769LH |  | 7.04 | 0 | Yes | Yes | Yes |

To investigate the possible role of NAMPT in formation of MN adducts, we assessed whether the NAMPT inhibitor GNE-617 (30) prevented formation of NB-3-MN in THP-1 cells, with Vacor as a positive control (9). Consistent with the role of NAMPT in NAD salvage pathways (30), NAMPT inhibition strongly reduced NAD levels and as expected Vacor-MN formation was prevented. However, we found that NAMPT was not required for NB-3-MN formation in THP-1 cells (Supplemental Figure S2), suggesting that an additional enzymatic mechanism exists for MN adduct formation in human cells. There was no correlation of NB-3-MN or Vacor-MN formation with cADPR or cADPR/NAD in THP-1 cells (data not shown), consistent with the lack of SARM1 expression in this cell line (31).

### Pyridine-containing drugs form MN adducts in vitro

Given our evidence that multiple enzymes had sufficient substrate promiscuity to generate MN adducts from BEI, we wondered if this phenomenon might extend to other pyridine-containing compounds. Using our THP-1 assay as a convenient in vitro tool, we screened pyridine-containing approved drugs and were intrigued to detect adducts formed by several compounds (Table 2). We observed MN adduct formation by chloroquine and indinavir, confirming that there is potential for MN adduct formation with other pyridine-containing compounds beyond BEI and known neurotoxins. We also tested two tuberculosis drugs, isoniazid and ethionamide, which have been previously shown to form drug-AD adducts that inhibit the *M. tuberculosis* InhA enzyme (32, 33). MN adducts were not detected for either of the tuberculosis base-exchange inhibitors, but the isoniazid-AD adduct was detected in THP-1 cells.

**Table 2.**
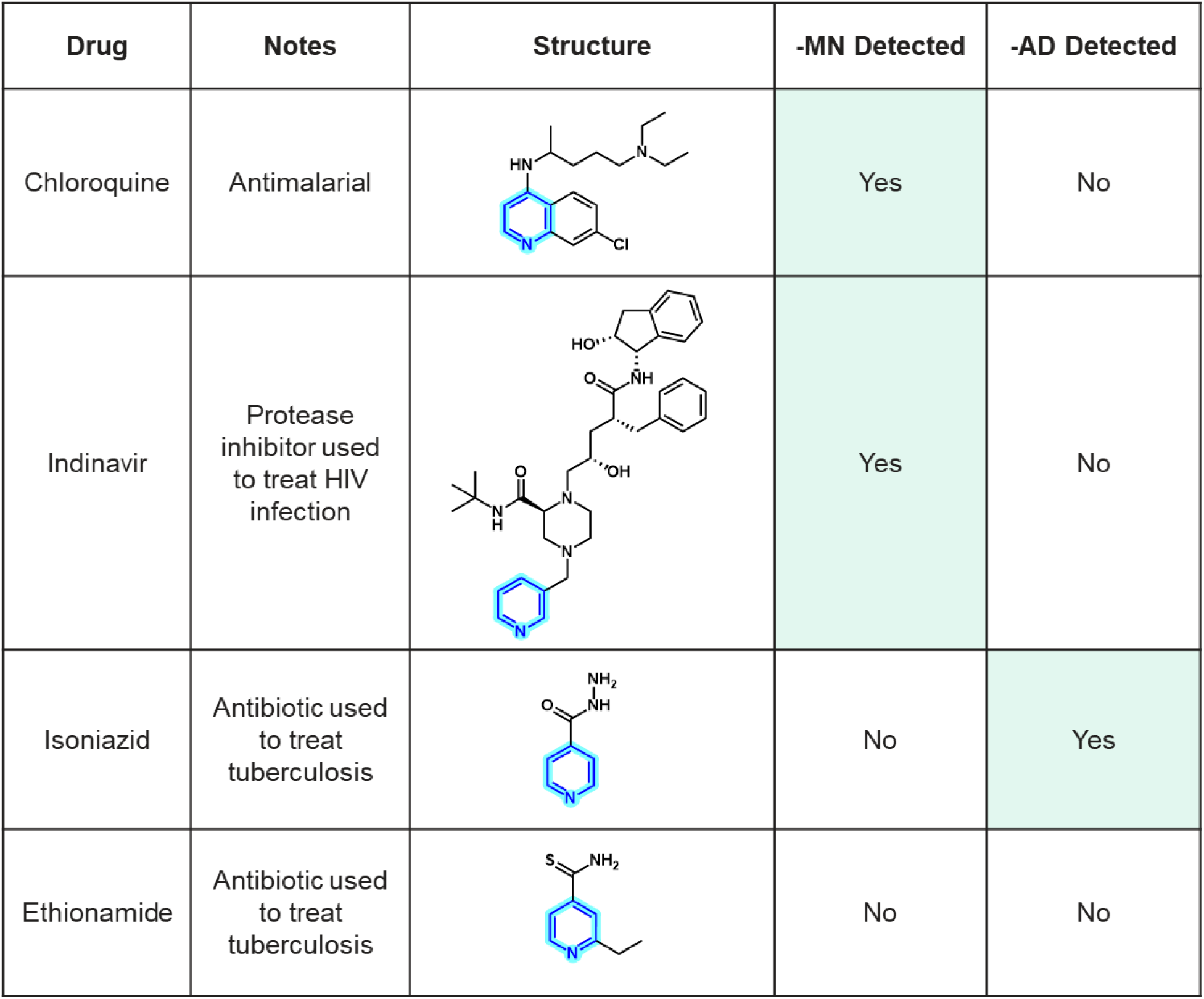
Pyridine-containingapproved drugs also form mononucleotideadducts in human cells.

| Drug | Notes | Structure | -MN Detected | -AD Detected |
| --- | --- | --- | --- | --- |
| Chloroquine | Antimalarial |  | Yes | No |
| Indinavir | Protease inhibitor used to treat HIV infection |  | Yes | No |
| Isoniazid | Antibiotic used to treat tuberculosis |  | No | Yes |
| Ethionamide | Antibiotic used to treat tuberculosis |  | No | No |

### Noncompetitive inhibitors do not activate SARM1

Although all our data suggested a correlation between BEI-MN adduct formation and SARM1 activation, it was still formally possible that the activation effect we had observed might extend to all SARM1 inhibitors regardless of mechanism of action. It was recently reported that low doses of BEI exacerbated the cytotoxic effects of Vacor in vitro (34–36), consistent with the SARM1 activating effects we had observed in vivo, while the allosteric covalent SARM1 inhibitor EV-99 lacked cytotoxic effects (36). We therefore replicated this assay with BEI and used it to assess representatives of two different non-BEI mechanisms. For this experiment, we selected MY-9B, which had been previously shown to inhibit SARM1 through allosteric covalent binding to C311 in the ARM domain (37). We also characterized a non-covalent inhibitor A1857PZ that we found to inhibit SARM1 through a noncompetitive mechanism, as evidenced by Michaelis-Menten enzyme kinetics studies showing decreased Vmax with no change in NAD Km in the presence of inhibitor and intersecting rather than parallel Lineweaver-Burk plots with and without inhibitor (Figure 6A). As previously reported, we found that the BEI NB-3 exacerbated cytotoxicity of Vacor at low doses and switched to protection from cytotoxicity at higher doses (Figure 6B). However, both non-BEI SARM1 inhibitors only exhibited protection from Vacor-induced cytotoxicity, with no exacerbation at lower doses.

**Figure 6.**
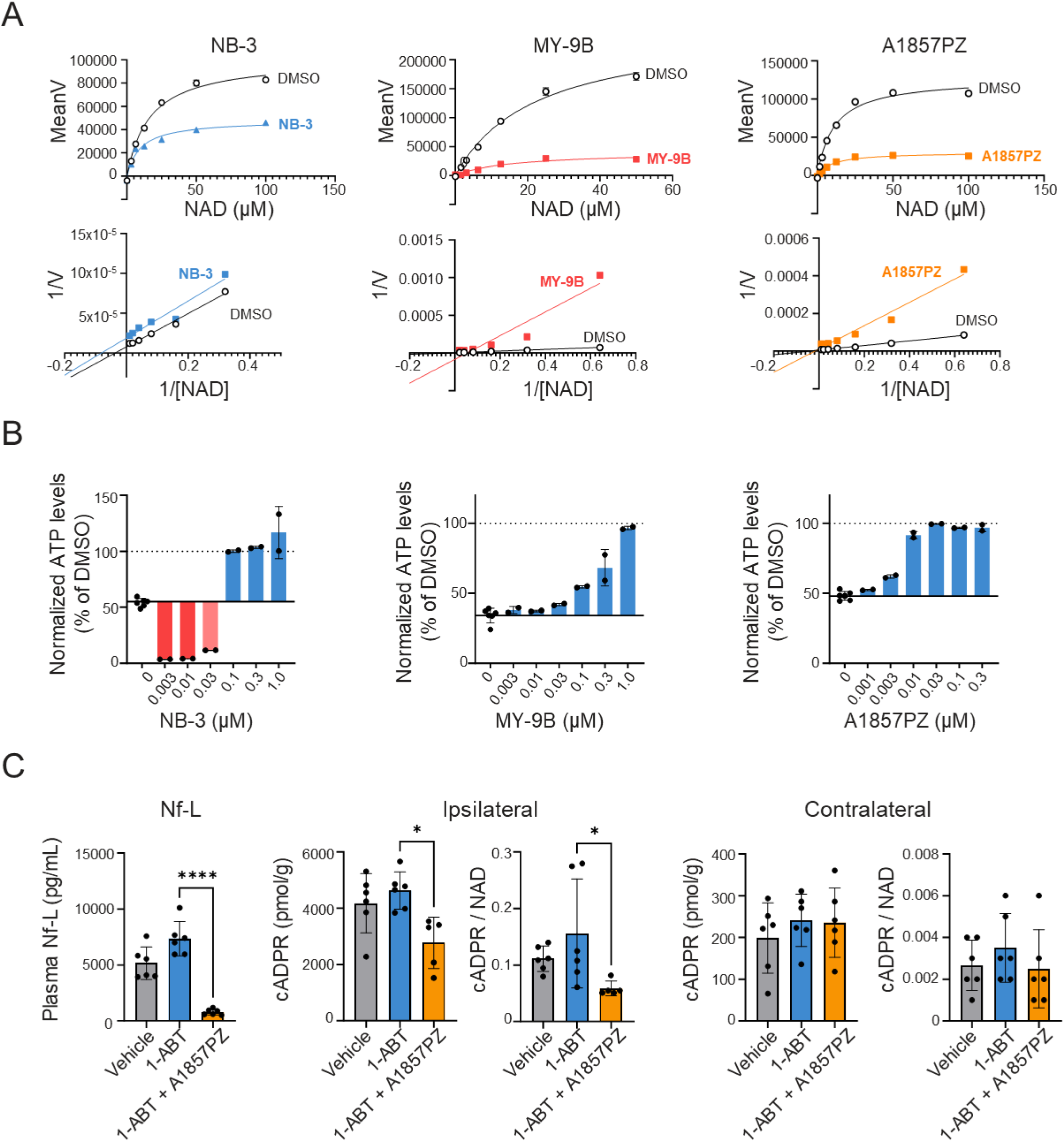
Noncompetitive inhibitors do not activate SARM1. **(A)** *In vitro* SARM1 activity assay using PC6 as a fluorescent readout for recombinant full-length SARM1 activity. Inhibitor effects on SARM1 NAD kinetics were determined by carrying out an NAD dose response at sub-saturating inhibitor concentration (IC50 or IC50*2). MoA was designated using Vmax and Km changes from Michaelis-Menten analysis (top) and Lineweaver-Burk plots (bottom). NB-3 (blue) represents a known BEI compound (uncompetitive); MY-9B (red) and A1857PZ represent non-BEI, noncompetitive compounds. **(B)** Cytotoxicity assay measuring SH-SY5Y cell viability in response to Vacor challenge (12.5 uM) in the presence of the BEI NB-3 or non-BEI compounds (MY9B, A185YPZ). **(C)** Plasma Nf-L and cADPR/NAD measurements in animals dosed with non-BEI compound A1857PZ. Results are shown as mean ± SEM. N ≥ 5 animals per condition. *, p<0.05; ****, p<0.0001 calculated using One-way ANOVA with Tukey’s multiple comparison test.

We next sought to confirm these results in vivo. Although MY-9B inhibited SARM1 in cells, its pharmacokinetic properties were not suitable for in vivo dosing (data not shown). However, we found that dosing the inhibitor A1857PZ 30 minutes after 50 mg/kg of the pan-cytochrome P450 inhibitor 1-aminobenzyltriazole (1-ABT) enabled us to achieve sufficient compound exposure in vivo (Supplemental Figure S3). In the SNA model, we observed neuroprotection (suppression of Nf-L) 20 h post-axotomy in animals dosed with 30 mg/kg A1857PZ 8 h after axotomy, along with reduction of cADPR and cADPR/NAD in the ipsilateral sciatic nerve consistent with SARM1 inhibition. No elevation in cADPR or cADPR/NAD was detected in contralateral sciatic nerve in dosed animals (Figure 6C).

## Discussion

We present herein examples of compounds that form a SARM1 inhibitor adduct through a base exchange mechanism (BEI) and paradoxically also activate SARM1 in uninjured nerves and in healthy, naïve animals. While this manuscript was in preparation, several other groups reported exacerbated nerve damage from low doses of BEI in vivo in acute nerve injury and disease models (34, 36, 38); however, to our knowledge this is the first report of SARM1 activation by BEI in healthy animals and uninjured nerves. By directly measuring SARM1’s product cADPR and the product to substrate ratio (cADPR/NAD) as biomarkers of enzymatic activity, we observed that BEI induced elevated cADPR and cADPR/NAD in healthy sciatic nerve. Catalytically inactive SARM1 KI mice were protected from elevated cADPR and cADPR/NAD after BEI dosing, confirming that this effect was due to SARM1 activation. Moreover, based on our discovery of BEI-MN adducts in vivo we propose a possible mechanism for the SARM1 activating effect we and others have observed.

It has previously been noted that cooperativity between subunits in multimeric enzymes can paradoxically lead to increased activity when substrate analog inhibitors bind at substoichiometric levels (39). BEI-adenine dinucleotide (BEI-AD) adducts could be considered substrate analogs; however we have seen no evidence that SARM1 subunits display cooperativity upon NAD binding. In our NAD Km determinations, the rate of SARM1 enzymatic activity in response to varying concentrations of NAD exhibits classic Michaelis-Menten curves (Figure 6A) rather than the distinctive sigmoidal shape that would signify cooperativity.

Instead, we propose that BEI form adducts that act as NMN analogs, activating SARM1 in a manner analogous to the known mechanisms of the SARM1 activator and former rodenticide Vacor (9) and the neurotoxic agent 3-acetylpyridine (3-AP) (8). Like Vacor and 3-AP, BEI also formed mononucleotide (MN) adducts, and BEI chemotypes that activated SARM1 in vivo formed BEI-MN adducts in sciatic nerve. We also noted that the time course of BEI-MN adduct formation matched the time course of cADPR elevation in sciatic nerve (Figure 4D, E). While our results do not enable us to conclude on causality, BEI-MN adducts can be considered a predictive biomarker for SARM1 activation.

By dosing the BEI NB-3 in SARM1 KI mice, we determined that multiple enzymes generate MN adducts in vivo, and the enzymes required for adduct formation vary in different tissues. We speculate that some of the many enzymes that use nicotinamide or its derivatives as substrates may indiscriminately act on pyridine-containing compounds such as BEI. The enzymes responsible for forming BEI-MN adducts may also vary by chemotype. In support of this idea, we confirmed a literature report that inhibiting NAMPT prevented formation of Vacor-MN (9), but NAMPT inhibition did not prevent formation of MN adducts for BEI.

Taken together, these results suggest that MN adduct formation may be highly context-dependent, and the ability to detect these metabolites may depend on the tissue being analyzed and cannot be generalized from a single BEI chemotype. These idiosyncrasies could present challenges for detecting MN adducts in preclinical metabolite identification studies. We developed an assay to detect adducts in the THP-1 human-derived cell line, confirming that adduct formation is not a rodent-specific phenomenon. While our cellular assay can be used to aid in vitro identification of compounds with a propensity to form MN adducts, it must be cautioned that the enzymatic proteome in THP-1 cells may not fully recapitulate potential for MN adduct formation in vivo in different tissue compartments. Similarly, transient SARM1 activation may be difficult to detect in preclinical studies, increasing the likelihood that compounds with these effects could advance to clinical testing with no apparent red flags.

Ingestion of Vacor in humans rapidly leads to neuropathy and new-onset diabetes (40, 41). Unlike Vacor, the SARM1-activating effect of BEI may be self-limiting due to formation of the inhibitor AD adduct by activated SARM1. We theorize that the two species of adducts, AD and MN, have opposite and competing effects, so the net effect of activation or inhibition depends on the concentration ratio of MN to AD adduct, and the relative potency of each. Consistent with this model, we observed that MN formation predominated at low plasma concentrations of BEI, coinciding with net activation in vivo. At high doses of BEI the accumulation of AD adduct initially resulted in net inhibition of SARM1, but we still observed net activation during washout periods after oral dosing at 30 mg/kg (Figure 4) or 100 mg/kg (Figure 5). Providing further support, it was recently reported that at low doses BEI (≤ 3 mg/kg) acutely exacerbated nerve damage in vivo in axotomy, low dose Vacor, and EAE models, while higher doses of BEI (≥ 30 mg/kg) were protective (34, 36). The long-term effects of BEI dosing are currently unknown, but we speculate that chronic oral dosing of a BEI could result in daily periods of net SARM1 activation during trough periods, and perhaps prolonged activation if a dose were accidentally missed. It is unclear what, if any, level of repeated SARM1 activation can be tolerated without induction of nerve damage or exacerbation of neurodegenerative disease.

Sporadic reports have suggested associations of pyridine-containing drugs with neuropathy after dosing in humans (42–47). Two drugs used to treat tuberculosis, isoniazid and ethionamide, are known to act by a base exchange mechanism, forming AD adducts that inhibit the *M. tuberculosis* InhA enzyme (32, 33). Both drugs are associated with development of peripheral neuropathy in patients undergoing treatment for tuberculosis, and Wallerian degeneration of peripheral nerves was observed in human post-mortem studies after isoniazid treatment (48, 49). We detected the AD adduct only for isoniazid in our THP-1 assay; the known AD adduct of ethionamide was not found, perhaps due to a requirement for *M. tuberculosis* enzymatic pathways not present in THP-1 cells. Similarly, we cannot exclude the possibility that MN adducts may form in the presence of *M. tuberculosis*.

However, two other pyridine-containing drugs, chloroquine and indinavir, did form MN adducts in the THP-1 assay. It is important not to overinterpret our in vitro assay results, but this evidence underscores the promiscuity of nicotinamide-related substrate recognition for adduct formation. Indinavir, a drug used to treat human immunodeficiency virus (HIV), is associated with sensory neuropathy in patients receiving the drug and causes neurite degeneration in vitro in HIV-infected dorsal root ganglion neurons (47, 50). Chloroquine is an antimalarial that has been associated with sensorimotor polyneuropathy and/or retinopathy after long-term treatment (51–55). The potential role of environmental exposures and low-level SARM1 activation in some forms of neuropathy is worthy of future study.

Neurotoxicity ranks as the third leading cause of approved drugs being pulled from the market, accounting for 12.5% of withdrawn drugs worldwide in the period from 1950-2016 (56). There is a continuing need to identify structural alerts, neurotoxic mechanisms, and methods to assess risk preclinically, and our work contributes to this understanding around pyridine-containing compounds. The observation that BEI can activate SARM1 in healthy animals and uninjured nerves may have important implications if clinical studies are conducted with BEI. Even if acute toxicity is not observed, the cumulative effect of repeated cycles of SARM1 activation may mask any therapeutic benefit of SARM1 inhibition in patients with neurodegenerative disease, perhaps causing loss of confidence in SARM1 as a drug target and missed opportunities to treat diseases with high unmet need.

Importantly, the effects we describe here do not apply to SARM1 inhibitors with differentiated mechanisms of action. In contrast to BEI, both noncompetitive and covalent allosteric SARM1 inhibitors did not exacerbate Vacor cytotoxicity in vitro; the absence of cytotoxicity with non-BEI mechanisms is in agreement with a recent result with the covalent allosteric inhibitor EV-99 (36). In addition, a noncompetitive SARM1 inhibitor dosed in vivo protected against nerve damage after sciatic nerve axotomy with no evidence of SARM1 activation in the uninjured nerve. Thus, while our results point to potential risks of the BEI mechanism, we also show that alternative modes of SARM1 inhibition may avoid these risks and provide a more promising path forward.

## Methods

### Reagents

Approved drugs were purchased from TargetMol except indinavir, which was purchased from Sigma. All other compounds were synthesized at Lundbeck La Jolla Research Center (San Diego). Compounds were solubilized in 100% dimethylsulfoxide (DMSO) for in vitro experiments.

### Animals, animal welfare, SARM1 KI mice

All animal experiments were approved by the Charles River and Accelerator Development Lab (CRADL) Institutional Animal Care and Use Committee. Animals were housed in ventilated cages and maintained on a 12-hour / 12-hour light / dark cycle and had free access to food and water through the duration of the experiments. Male Sprague Dawley Rats from Charles River Laboratories and male C57Bl/6J from Jackson Laboratories mice were 5-7 weeks old at the time of experiments. SARM1^E642A /E642A^ knock-in mice were generated on a BALB/cAnNTac background by Taconic Biosciences GmbH via CRISP/Cas9 gene editing to mutate the catalytic glutamate 642 to alanine, rendering SARM1 NADase activity non-functional. Briefly, a guide RNA targeting exon 8 of the mouse *Sarm1* gene was selected for its position and low number of predicted off-targets. Founder mice were identified by PCR and bred with WT BALB/cAnNTac mice to generate F1 Heterozygous mice. SARM1^E642A/WT^ heterozygous mice were bred to generate mixed sex littermate cohorts that were used for experiments at 10-20 weeks old.

### Formulation and dosing of SARM1 inhibitors

NB-2, NB-3, A1715CG, A1767FN, A1767FO, A1713DK, and A1769LH were formulated in 2.5% DMSO, 10% Cremophor EL, and 85% sterile filtered pH 7.0 water with 5% dextrose and administered by oral gavage (p.o.) at a volume of 5 mL/kg (rats) or 10 mL/kg (mice). A1857PZ was formulated in 0.5% hydroxypropyl methylcellose (HPMC) and administered p.o. at 10 mL/kg. The pan-cytochrome P450 inhibitor, 1-aminobenzyltriazole (1-ABT, Sigma Aldrich) was dissolved in 0.5% methylcellulose (MC, Sigma Aldrich) and administered orally at 2 mL/kg 30 minutes before A1857PZ to improve in vivo stability. SARM1 inhibitors were administered to naïve mice 0 or 8 hours after sciatic nerve axotomy (SNA) and blood samples were collected by submandibular or submental bleed and placed into tubes containing K_2_EDTA for NfL and compound concentration analysis. For PK studies, blood samples were collected from the saphenous vein into K_2_EDTA tubes at 0.083, 0.25, 0.5, 1, 2, 4, 6, 8, 12 and 24 hours after administration.

### Sciatic nerve axotomy

Sciatic nerve axotomy (SNA) was performed in mice and rats using similar methods appropriately scaled to the size of the animal. Animals were anesthetized with isoflurane (3-4 %) in 100% oxygen and the surgical site on the hind limb was shaved and disinfected by 3 alternating scrubs of providone-iodine solution and 70% isopropyl alcohol. Animals were administered either carprofen (rats, 5 mg/kg, s.c.) or Ethiqa-XR (3.25 mg/kg for mice, 0.65 mg/kg for rats, s.c.) to manage post-operative pain. At the mid-thigh level an incision was made and the sciatic nerve exposed by blunt dissection. The sciatic nerve was cut distal to the sciatic notch and a ∼1 mm section was removed to prevent anastomosis of the cut ends. The surgical site was closed using wound clips and animals were returned to cages containing hydrogels (ClearH2O) and chow on the bottom of the cage to aid with hydration and ensure access to food. Between 12-72 hours after SNA, blood was collected by cardiac puncture into K_2_EDTA tubes for NfL and tissues were collected and frozen on liquid nitrogen and stored at -80 °C until analysis.

### Neurofilament light chain assay

Plasma neurofilament light chain was measured using the UmanDiagnostics NF-light Serum ELISA Assay (Quanterix 20-8002), according to manufacturer instructions.

### THP-1 cell treatments

THP-1 cells were plated in a twelve-well plate at a density of 1.1 x 10^6^ cells per well and treated with Vacor (50 μM) or NB-3 (100 μM) with or without the presence of NAMPT inhibition (GNE-617; 1 μM) for 24 hours. Cell pellets were harvested and analyzed as described below.

### cADPR, NAD, -MN and -AD adduct measurement in cells and tissues

Endogenous metabolites and drug adducts concentration in cells, sciatic nerve and liver was determined by ultra-performance liquid chromatography coupled to mass spectrometry (UPLC/MS). Biological material homogenization slightly differed between the three matrices, but the rest of the extraction and LC/MS analysis was common.

Cell pellets were harvested by collecting the THP-1 cells and growth media into a safelock Eppendorf Tube. Cells were pelleted (300 x g for 5 minutes at 4 °C) and the media was removed. Afterwards, cell pellets were washed with 500 mL of PBS. Finally, cells were spun down to remove the PBS (300 x g for 5 minutes at 4 °C) and snap frozen under liquid nitrogen for 1 min and stored at -80 °C until analyzed. 500 mL of pre-cooled (-20 °C) acetonitrile/methanol/water 2:2:1 (by vol.) was added to the extracts, containing 50 pmol of internal standard compounds (8-Br-cADPR, NAD-d4 and NMN-d4). Samples were vortexed for 30 seconds and incubated for 1 hour at -80 °C. Rest of the extraction protocol was common to tissues.

For sciatic nerve samples, 100 mL of pre-cooled (-20 °C) methanol/water 4:1 (by vol.) was added to the extracts, containing 50 pmol of internal standard compounds (8-Br-cADPR, NAD-d4 and NMN-d4). Tissues were homogenized for 30 seconds at 30 Hz twice in a tissue lyser (Tissue Lyser II, Qiagen, Venlo, Netherlands) with one stainless steel bead (5 mm diameter). Samples were vortexed and kept on ice between both homogenization cycles. After homogenization, 400 mL of pre-cooled (-20 °C) methanol/water 4:1 (by vol.) was added to the extracts without containing any standard. Samples were vortexed for 30 seconds, sonicated in an icy bath for 15 min and incubated for 1 hour at -20 °C. Liver homogenization followed the same protocol but adding the 500 mL of pre-cooled (-20 °C) methanol/water 4:1 (by vol.) at once (containing 50 pmol of each internal standard) and running a single mechanical homogenization cycle. Rest of the extraction protocol was common to cells.

Cells or tissue homogenates were centrifuged at 21,000 x g for 15 min at 4 °C and supernatants were transferred to another tube. Solvent was evaporated under a nitrogen stream. 70 mL of ice-cold acetonitrile/water 1:1 (v/v) containing 10 mM ammonium formate were added. Samples were vortexed for 30 seconds and sonicated in an icy bath for 10 minutes. Finally, metabolite extracts were centrifuged at 21,000 x g for 5 min at 4 °C to remove the non-soluble debris and 60 mL of the supernatants were transferred to a LC vial containing a borosilicated glass insert for UPLC/MS analysis.

Sample extracts (5 mL) were injected onto an Agilent 1290 UPLC system equipped with a G7120A pump, a G7129B autosampler and a G7116B column manager (Agilent Technologies, Santa Clara, CA). Chromatographic separation was achieved using an Acquity UPLC BEH Amide Column (2.1 x 100 mm, 1.7 mm particle size, 130 Ǻ) coupled to an Acquity UPLC BEH Amide Column Guard (2.1 x 5 mm, 1.7 mm particle size, 130 Ǻ) (Waters Corporation, Milford, MA). Mobile phase A was composed of water/acetonitrile 95:5 (v/v) and mobile phase B was composed of acetonitrile/water 95:5 (v/v) with 0.1% formic acid and 10 mM ammonium formate added to both mobile phases. The gradient started with 100% B for 2 min before being decreased linearly to 65% B over 12 min. Then, B solvent was decreased linearly to 40% over 3 minutes. Afterwards, solvent B was kept at 40% for 1 min, before switching to the initial conditions in 0.1 min. The system was allowed to equilibrate for 3.4 minutes before the next sample injection. Flow rate was kept at 0.4 mL/min and the column temperature was 27 °C. Analytes were quantified using a 6475 triple quadrupole mass spectrometer equipped with an electrospray Jet Stream source (Agilent Technologies) operated in dynamic multiple reaction monitoring (dMRM) mode. The following parameters were kept constant for all transitions: Fragmentor=100, Cell Accelerator Voltage=4, Polarity=Positive. Total cycle time was 650 ms. Source parameters were kept as follows: Dry Gas Temperature=350 °C, Dry Gas Flow=11 L/min, Sheath Gas Temperature=350 °C, Sheath Gas Flow=12 L/min, Nebulizer= 45 psi, Noozle voltage= 1500 V (positive) and Capillary= 3500 V (positive).

The quantitative and qualitative transitions for each metabolite, internal standard and drug with commercially available analytical standard (cADPR, NAD, NMN, 8-Br-cADPR, NAD-d4 and NMN-d4, all BEI and pyridine-containing approved drugs) were optimized using the Optimizer software (Agilent Technologies). The parent mass ([M+H]+ ion) of the -MN and -AD adducts formed with the BEI and the pyridine drugs was calculated by analogy with the NMN and NAD structures, exchanging nicotinamide by the drug structure (see table below). MS/MS transitions (1 quantifier and 2 qualifiers) were predicted leveraging the known fragmentation pattern of NAD and NMN analytical standards (and their known deuterated isotopologues). The quantifier transition for the -MN adduct was the [M+H]+ ion formed from the free drug; while the qualifiers were the main transition of the free drug and 97.0, which is known to be specific for the ribose structure. The quantifier transition for the -AD adduct was 136.1, originated from adenine. The qualifiers for the -AD adduct formation were 524.1 (from the ADP-ribose part of the molecules) and the [M+H]+ ion coming from the free drug. ion This prediction was confirmed in the bibliography for Vacor and isoniazid forming adducts, using synthesized standards and high-resolution mass spectrometry (33, 57). All the identified adducts following this procedure were subjected to further evaluation to rule out possible false positives. Only peaks meeting the following criteria were considered hits: a) at least 3 transitions were found in the drug-treated samples, but not in vehicle-treated controls. b) Relative abundance of fragments resembles their equivalent in NAD and NMN. c) Chromatographic retention time also resembles the endogenous molecules (free drug<-MN<-AD). d) -MN and -AD adduct parent ion was fragmented at 3 different collision energies (10, 20, 40 eV) and the same fragments were found in all of them (with different abundance) in the drug-treated samples, but not the controls.

Metabolites were quantified using an isotope dilution method by comparing the integrated area of each endogenous analyte to the area of its corresponding isotopic labelled analogue using the MassHunter Quantitative Software (Agilent Technologies). NMN-d4 and NAD-d4 were used as internal standards for the BEI and pyridine drug -MN and -AD forming adducts, respectively, although the data is not fully quantitative since we do not have the analytical standard for each compound.

Calibration curves using analytical standards were run before the experiments to evaluate lower limit of detection, linearity and dynamic range of the method. Finally, data was normalized to the initial number of cells or tissue weight.

| Metabolite | Internal Standard | Parent Ion [M+H] | Quantifier Transition (CE in eV) | Qualifier Transition(s) (CE in eV) |
| --- | --- | --- | --- | --- |
| **cADPR** | *8-Br-cADPR* | 542.1 | 542.1→ 136.0 (49) | 542.1→ 348.1 (25) |
| **NAD** | *NAD-d4* | 664.1 | 664.1→ 136.0 (53) | 664.1→ 428.2 (21) |
| **NB-2-MN** | *NMN-d4* | 460.1 | 460.1→ 248.1 (25) | 460.1→ 121.1 (45), 97.0 (45) |
| **NB-2-AD** | *NAD-d4* | 789.2 | 789.2→ 136.1 (53) | 789.2→ 524.1 (21), 248.1 (50) |
| **NB-3-MN** | *NMN-d4* | 474.1 | 474.1→ 262.1 (25) | 474.1→ 135.1 (45), 97.0 (45) |
| **NB-3-AD** | *NAD-d4* | 803.2 | 803.2→ 136.1 (53) | 803.2→ 524.1 (21), 262 (50) |
| **A1715CG-MN** | *NMN-d4* | 439.1 | 439.1→ 227.1 (25) | 439.1→ 171.0 (45), 97.0 (45) |
| **A1715CG-AD** | *NAD-d4* | 768.1 | 768.1→ 136.1 (53) | 768.1→ 524.1 (21), 227.1 (50) |
| **A1767FN-MN** | *NMN-d4* | 422.1 | 422.1→ 210.1 (25) | 422.1→ 182.0 (45), 97.0 (45) |
| **A1767FN-AD** | *NAD-d4* | 751.2 | 751.2→ 136.1 (53) | 751.2→ 524.1 (21), 210.1 (50) |
| **A1767FO-MN** | *NMN-d4* | 423.1 | 423.1→ 211.1 (25) | 423.1→ 183.0 (45), 97.0 (45) |
| **A1767FO-AD** | *NAD-d4* | 752.2 | 752.2→ 136.1 (53) | 752.2→ 524.1 (21), 211.1 (50) |
| **A1713DK-MN** | *NMN-d4* | 562.2 | 562.2→ 350.1 (25) | 562.2→ 197.0 (45), 97.0 (45) |
| **A1713DK-AD** | *NAD-d4* | 891.2 | 891.2→ 136.1 (53) | 891.2→ 524.1 (21), 350.1 (50) |
| **A1769LH-MN** | *NMN-d4* | 474.1 | 474.1→ 262.1 (25) | 474.1→ 135.1 (45), 97.0 (45) |
| **A1769LH-AD** | *NAD-d4* | 803.2 | 803.2→ 136.1 (53) | 803.2→ 524.1 (21), 262 (50) |
| **Isoniazid-MN** | *NMN-d4* | 350.1 | 350.1→ 138.1 (25) | 350.1→ 110.0 (45), 97.0 (45) |
| **Isoniazid-AD** | *NAD-d4* | 679.2 | 679.2→ 136.1 (53) | 679.2→ 524.1 (21), 138.1 (50) |
| **Ethionamide-MN** | *NMN-d4* | 379.1 | 379.1→ 167.1 (25) | 379.1→ 107.0 (45), 97.0 (45) |
| **Ethionamide-AD** | *NAD-d4* | 708.2 | 708.2→ 136.1 (53) | 708.2→ 524.1 (21), 167.1 (50) |
| **Indinavir-MN** | *NMN-d4* | 826.4 | 826.4→ 614.4 (25) | 826.4→ 421.0 (45), 97.0 (45) |
| **Indinavir-AD** | *NAD-d4* | 1155.5 | 1155.5→ 136.1 (53) | 1155.5→ 524.1 (21), 614.4 (50) |
| **Chloroquine-MN** | *NMN-d4* | 532.2 | 532.2→ 320.2 (25) | 532.2→ 247 (45), 97.0 (45) |
| **Chloroquine-AD** | *NAD-d4* | 861.3 | 861.3→ 136.1 (53) | 861.3→ 542.1 (21), 320.2 (50) |
| **8-Br-cADPR** | - | 620.0 | 620.0→ 214.0 (33) | 620.0→ 97.0 (53) |
| **NAD-d4** | - | 668.1 | 668.1 → 136.0 (53) | 668.1 → 428.0 (30) |
| **NMN-d4** | - | 339.1 | 339.1 → 127.1 (10) | 339.1 → 97.1 (30) |
| **NMN** | - | 335.1 | 335.1 → 123.1 (10) | 335.1 → 97.0 (30) |

### Compound concentration in plasma, liver and sciatic nerves

Compound concentration in plasma, liver and sciatic nerves was determined by ultra-performance liquid chromatography coupled to mass spectrometry (UPLC/MS). Blood was collected into K2-EDTA containing tubes and kept on ice until processed. Blood samples were centrifuged at 9,000 x g for 3 minutes at 4 °C and the plasma supernatant was transferred to another tube and frozen on dry ice and stored at -80 °C until analyzed. The calibration curve samples were prepared using plasma from naïve mice isolated following the same protocol as above and spiked with serial dilutions of the test article, ranging 1-50000 ng/mL. 25 mL of plasma were mixed with 100 mL of ice-cold acetonitrile, vortexed for 30 seconds, kept on ice for 15 minutes and centrifuged at 21,000 x g at 4 °C for 5 minutes. Supernatants were transferred to another tube. 20 mL of supernatant were mixed with 80 mL of water and transferred to a LC vial containing a borosilicate glass insert for UPLC/MS analysis. For liver, 3 volumes (mL) of ice-cold acetonitrile/water 3:1 (v/v) were added to frozen tissues by tissue weight (mg) for compound extraction. Afterwards, tissues were homogenized for one minute at 30 Hz in a tissue lyser (Tissue Lyser II, Qiagen, Venlo, Netherlands) with one stainless steel bead (5 mm diameter). Samples were incubated on ice for 30 minutes and then centrifuged at 2,400 g at 4 °C for 15 minutes and supernatants were transferred to another tube. The calibration curve samples were made using a liver homogenate from naïve mice (extracted using the same protocol as the test samples) spiked with serial dilutions of the test article, ranging 1-50000 ng/mL. 20 mL of supernatant were mixed with 80 mL of water and transferred to a LC vial containing a borosilicate glass insert for UPLC/MS analysis.

For sciatic nerve samples, 20 volumes (mL) of ice-cold methanol/water 4:1 (v/v) were added to frozen tissues by tissue weight (mg) for compound extraction. Afterwards, tissues were homogenized for 30 seconds at 30 Hz twice in a tissue lyser with one stainless steel bead (5 mm diameter). Samples were vortexed and kept on ice between both homogenization cycles. Samples were incubated for 60 minutes at -20 °C and then centrifuged at 21,000 x g at 4 °C for 15 minutes and supernatants were transferred to another tube. The calibration curve samples were made using a sciatic nerve homogenate from naïve mice (extracted using the same protocol as the test samples) spiked with serial dilutions of the test article, ranging 1-50000 ng/mL. 20 mL of supernatant were mixed with 80 mL of water and transferred to a LC vial containing a borosilicated glass insert for UPLC/MS analysis.

Sample extracts (10 mL) were injected onto an Agilent 1290 UPLC system equipped with a G7120A pump, a G7167B multisampler and a G1170A column manager (Agilent Technologies, Santa Clara, CA). Chromatographic separation was achieved using an Acquity UPLC BEH C18 Column (2.1 x 50 mm, 1.7 mm particle size, 130 Ǻ) coupled to an Acquity UPLC BEH C18 Column Guard (2.1 x 5 mm, 1.7 mm particle size, 130 Ǻ) (Waters Corporation, Milford, MA). Mobile phase A was composed of water/acetonitrile 95:5 (v/v) and mobile phase B was composed of acetonitrile/water 95:5 (v/v) with 0.1% formic acid added to both mobile phases. The gradient started with 0% B for 0.5 min before being increased linearly to 100% B over 4 min. Afterwards, solvent B was kept at 100% for 1 min, before switching to the initial conditions in 0.1 min. The system was allowed to equilibrate for 1.4 minutes before the next sample injection. Flow rate was kept at 0.6 mL/min and the column temperature was 50 °C.

Analytes were quantified using a 6470 triple quadrupole mass spectrometer equipped with an electrospray Jet Stream source (Agilent Technologies) operated in dynamic multiple reaction monitoring (dMRM) mode. The quantitative and qualitative transitions for each compound were optimized using the authentic standards in the Optimizer software (Agilent Technologies): NB-2 (Quant: 248.1➔ 121.1, CE=26 eV; Qual: 248.1➔ 80.1, CE= 45 eV); NB-3 (Quant: 262.1➔ 135.1, CE= 22 eV; Qual: 262.1➔ 80.1, CE= 45 eV); A1715CG (Quant: 227.1➔ 171, CE= 34 eV; Qual: 227.1➔ 199, CE= 26 eV). The following parameters were kept constant for all transitions: Fragmentor=140, Cell Accelerator Voltage=4, Polarity=Positive. Total cycle time was 300 ms. Source parameters were kept as follows: Dry Gas Temperature=350 °C, Dry Gas Flow=11 L/min, Sheath Gas Temperature=350 °C, Sheath Gas Flow=11 L/min, Nebulizer= 50 psi, Noozle voltage= 1500 V (positive) and Capillary= 3500 V (positive). Compounds concentrations were calculated by interpolating the integrated area under the curve with the calibration curves for each tissue prepared in the same matrix than the samples using the Masshunter Quantitative Analysis Software (Agilent Technologies).

### Mechanism of Action Analysis

SARM1 activity and NAD Km was assessed *in vitro* by preparing an NAD serial dilution at the IC50 for each inhibitor, 2X IC50, or with no inhibitor added (DMSO control). The reaction conditions contained 50mM HEPES pH 7.5, 10mM KCl, 1.5mM MgCl2, 40μM NMN, 20μM PC6 fluorogenic SARM1 activity reporter, described in (58), and 0.2 mg/mL EXPI cell lysate containing overexpressed human SARM1. Compound and NAD were pre-incubated with SARM1 lysate for 1.5 hours before adding the remaining reaction components. PC6 fluorescence was immediately assessed on a BioTek Synergy Neo2 plate reader (Ex 390/Em 520 nm) in kinetic mode over 30 minutes. To determine kinetic parameters, initial rates (mean slopes) were plotted versus [NAD] and the data was fit using a Michaelis Menten fit in GraphPad Prism 10. The 2X IC50 data was also graphed as a Lineweaver-Burk plot using GraphPad Prism 10. Mechanism of Inhibition (noncompetitive versus uncompetitive) was then determined using the two graphs.

### Vacor-induced cytotoxicity assay

Human neuroblastoma SH-SY5Y cells were plated at 15,000 cells/well in 96 well plates. The next day, cells were treated simultaneously with SARM1 inhibitor serial dilutions and 12.5 uM Vacor. After 24h, cytotoxicity was measured with CellTiter-Glo assay (Promega) according to manufacturer instructions.

---

**Figure S1.**
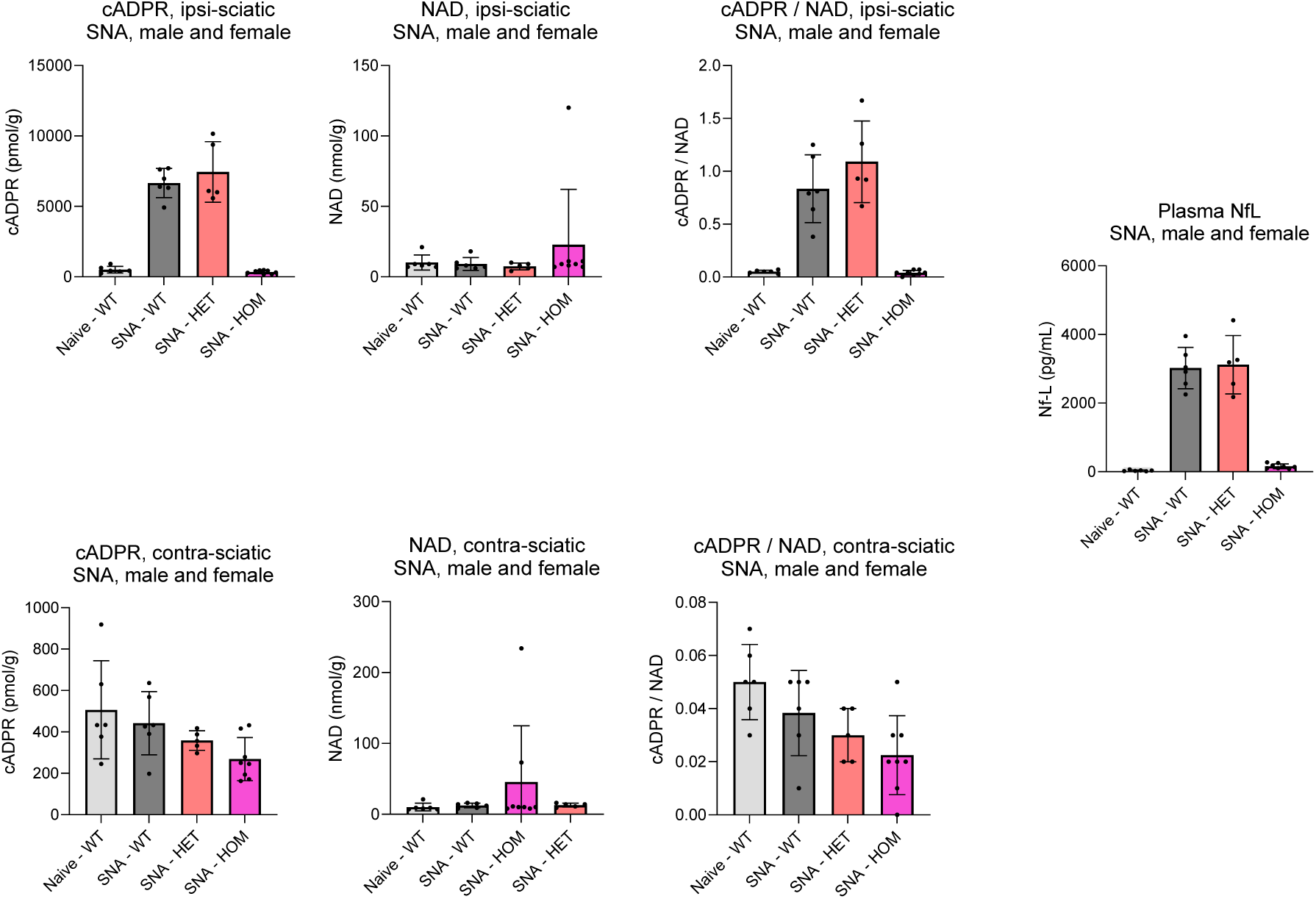
SARM1-E642A mutant is catalytically inactive *in vivo*. A mixed sex cohort of CRISPR knock-in (KI) mice with the catalytic glutamate of SARM1 mutated to alanine (E642A) were profiled in the SNA model, with measurement of plasma Nf-L and sciatic nerve cADPR and NAD.

**Figure S2.**
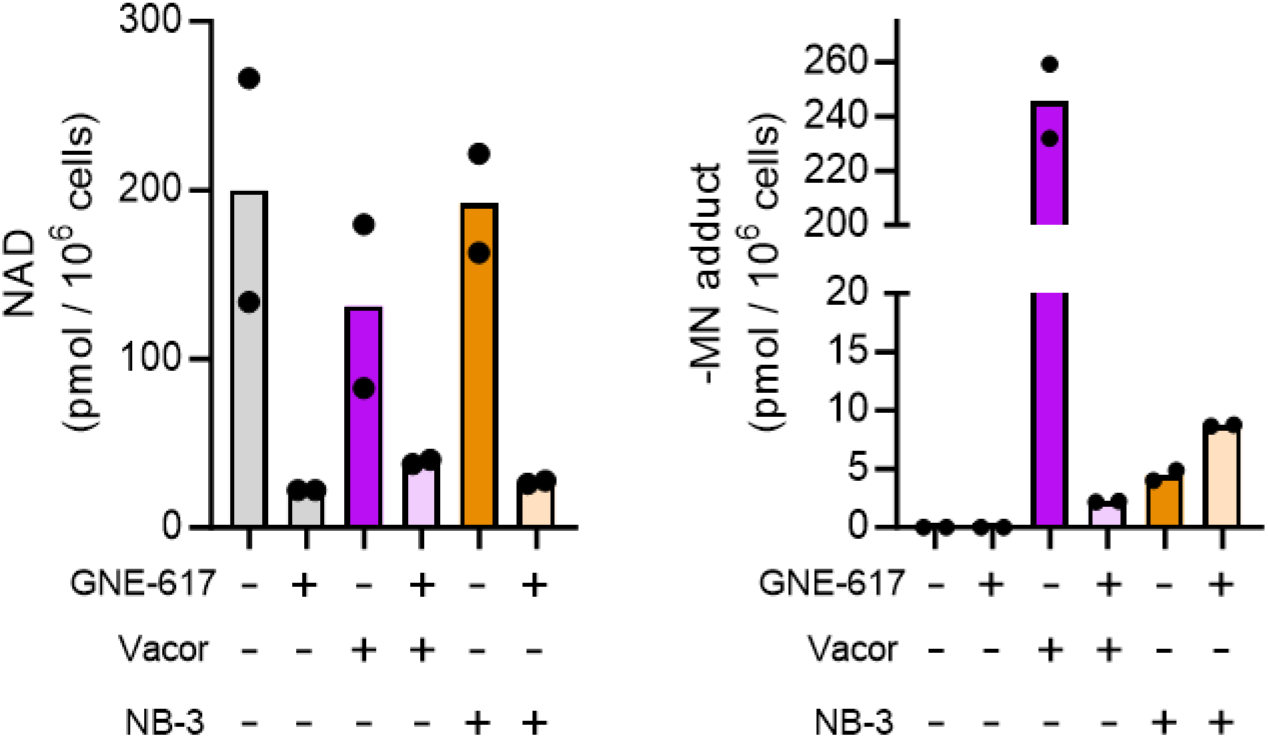
NAMPT inhibition blocks Vacor-MN formation but not NB-3-MN formation. NAD levels and MN adduct formation in THP-1 cells treated for 24 h with Vacor (50 μM) or NB-3 (100 μM) with or without the presence of NAMPT inhibition (GNE-617; 1 μM).

**Figure S3.**
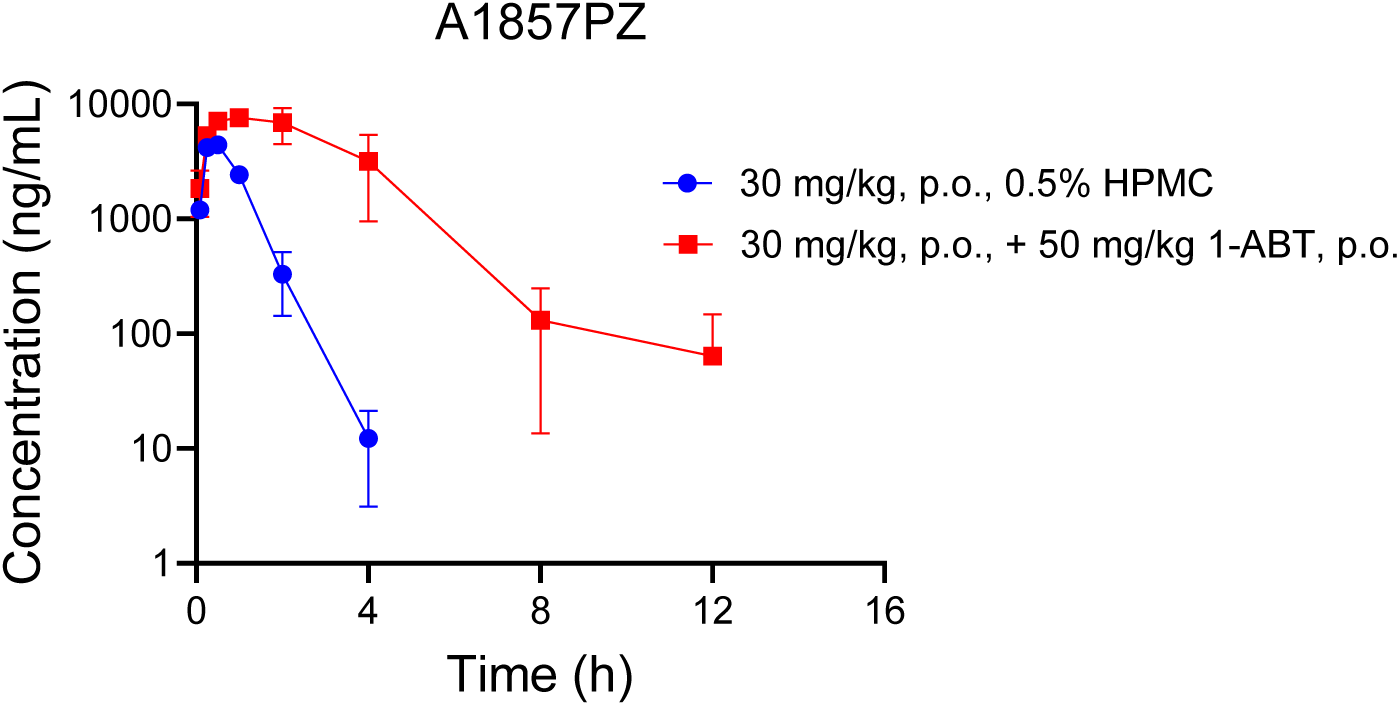
1-ABT treatment increases A1857PZ plasma concentrations *in vivo*. Time course of A1857PZ plasma concentrations following 30 mg/kg p.o. administration alone or 30 minutes after a 50 mg/kg p.o. dose of 1-ABT.

